# An obligate intracellular bacterial pathogen forms a direct, interkingdom membrane contact site

**DOI:** 10.1101/2023.06.05.543771

**Authors:** Yamilex Acevedo-Sánchez, Patrick J. Woida, Stephan Kraemer, Rebecca L. Lamason

## Abstract

Interorganelle communication regulates cellular homeostasis through the formation of tightly-associated membrane contact sites ^1–3^. Prior work has identified several ways that intracellular pathogens alter contacts between eukaryotic membranes ^4–6^, but there is no existing evidence for contact sites spanning eukaryotic and prokaryotic membranes. Here, using a combination of live-cell microscopy and transmission and focused-ion-beam scanning electron microscopy, we demonstrate that the intracellular bacterial pathogen *Rickettsia parkeri* forms a direct membrane contact site between its bacterial outer membrane and the rough endoplasmic reticulum (ER), with tethers that are approximately 55 nm apart. Depletion of the ER-specific tethers VAPA and VAPB reduced the frequency of rickettsia-ER contacts, suggesting these interactions mimic organelle-ER contacts. Overall, our findings illuminate a direct, interkingdom membrane contact site uniquely mediated by rickettsia that seems to mimic traditional host MCSs.

## INTRODUCTION

Eukaryotic organelles do not exist in isolation, and instead are directly tethered to one another through well-defined membrane contact sites (MCS) ^2, 3^. These sites bring organelle membranes within 10–80 nm of each other and rely on a cadre of protein tethers ^7, 8^. The roles of MCSs in cellular homeostasis are still emerging, with many impacting lipid metabolism and transport, organelle dynamics, autophagy, and apoptosis ^1^. At the center of these connections is the largest organelle, the endoplasmic reticulum (ER), which acts as a major cellular scaffold bridging multiple organelles to itself ^3^.

Recent research indicates that ER-mediated MCSs are susceptible to pathogen subversion ^9, 10^. Pathogens appear to either manipulate pre-existing MCSs ^4^ or establish new ones between the ER and the host-derived membrane that harbors the pathogen ^6^. However, there is currently no evidence of MCSs directly bridging eukaryotic and prokaryotic membranes. Direct interactions with the ER would likely be mediated by pathogens, such as *Rickettsia parkeri* (the tickborne bacterium causing spotted fever disease), *Listeria monocytogenes*, or *Shigella flexneri* (agents of foodborne diarrheal disease), who spend most of their intracellular life cycle freely inhabiting the host cytosol^11^.

Here, we used a combination of live-cell imaging, transmission electron microscopy (TEM), and focused-ion beam scanning electron microscopy (FIB-SEM) to investigate whether cytosolic bacterial pathogens form direct MCSs with the ER. We discovered that a subset of *R. parkeri* form their own MCSs with the host cell ER. The frequency of these contact sites dramatically increased for a mutant *R. parkeri* strain that is unable to form actin tails, but similar structures were never observed after infection with various strains of *L. monocytogenes* or *S. flexneri*. The ER typically enveloped more than half of the surface of *R. parkeri* with visible tethers spanning the bacterial and ER membranes. Consistent with this observation, knockdown of *VAPA* and *VAPB* expression decreases the frequency of cells with these tethered rickettsia-ER contact sites. Taken together, our work identifies a novel membrane contact site between the ER and a cytosolic bacterial pathogen, elucidating a direct, interkingdom interaction for the ER.

## RESULTS

### *R. parkeri* forms stable interactions with the ER

Cytosolic pathogens such as *S. flexneri, L. monocytogenes,* and *R. parkeri* are constantly exposed to the host cytoplasm, prompting us to screen them in search of direct MCSs between prokaryotic and eukaryotic membranes. To identify putative interactions between the ER and cytosolic pathogens, we built a stable human A549 cell line that expressed a fluorescent ER marker (mCherry-Sec61β; A549-mCh-Sec61β) ^12^ and infected them with either DasherGFP-expressing *L. monocytogenes*, GFP-expressing *S. flexneri*, or TagBFP-expressing *R. parkeri* (RpBFP). Using live-cell imaging we observed that neither *L. monocytogenes* nor *S. flexneri* showed obvious interactions with the ER **(Fig. S1a, b)**. Conversely, 3–4% of RpBFP colocalized with the ER at 28 and 48 h post-infection (hpi), and these interactions were subdivided into two distinctive patterns: vacuole-like and protrusive ER structures **(Fig. 1a, b)**. Protrusive ER structures were found extending into the nucleus or through the cytoplasm **(Fig. 1b)**. Vacuole-like ER structures around bacteria were more abundant and only found in the cytoplasm **(Fig. 1c)**. These colocalization patterns were independent of the fluorescent ER marker used, as we obtained similar results with another ER marker (BiP-mNeonGreen-KDEL) (**Fig. S2**) ^13^.

**Figure 1:**
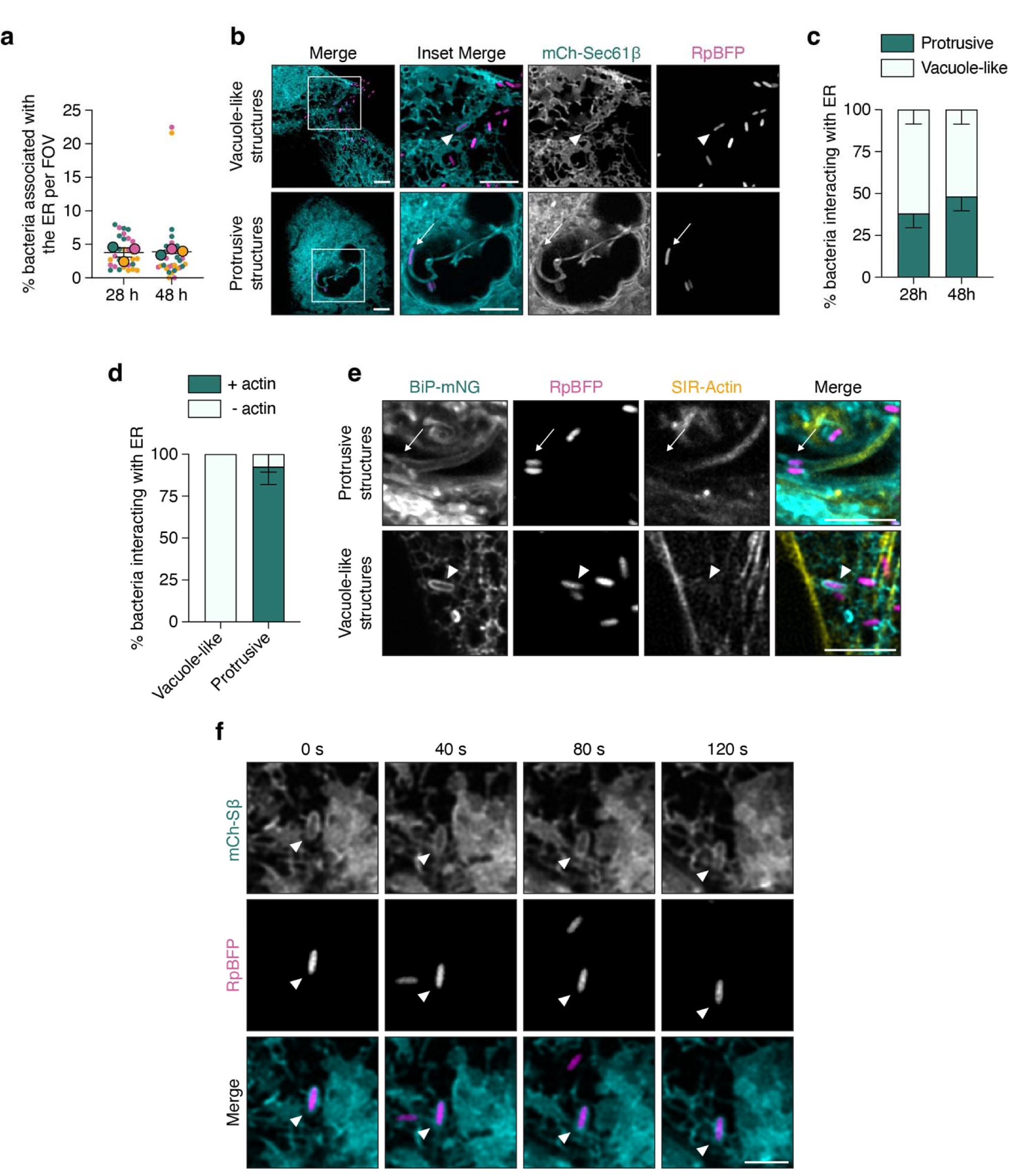
*R. parkeri* interacts with the ER. **a**, Percentage of bacteria interacting with all ER structures at 28 or 48 hpi, as defined in b. The mean ± SEM of 3 independent experiments (outlined circles) are superimposed over the raw data (unlined circles) representing the percentage per field of view (FOV) (> 2000 bacteria counted per independent experiment). **b**, Live-cell imaging snapshots at 28 hpi of RpBFP (WT *R. parkeri* expressing BFP, magenta) in vacuole-like (arrowhead) or protrusive (arrow) ER structures after infecting A549 cells expressing an ER marker (mCherry-Sec61β, cyan). Scale bar, 5µm; inset scale bar, 2.5µm. **c**, Relative frequency of bacteria interacting with the ER in B that are forming protrusive or vacuole-like structures at 28 or 48 hpi. Error bars indicate SEM from 3 independent experiments (> 80 bacteria counted per independent experiment). **d**, Relative frequency of bacteria interacting with the ER that display an actin tail (as in e). Data presented are mean ± SD. Approximately 10 fields of view were imaged per strain for each replicate (n = 2) and > 100 bacteria interacting with the ER were quantified. **e**, Live-cell imaging snapshots at 28 hpi of RpBFP (magenta) in vacuole-like (arrowhead) or protrusive (arrow) structures after infecting A549 cells expressing an ER marker (BiP-mNeonGreen-KDEL, cyan) and stained with SIR-actin (yellow). Scale bar, 5µm. **f**, Live-cell imaging snapshots showing vacuole-like ER structures around bacteria over time. Scale bar, 2.5µm.

During infection, a subpopulation of *R. parkeri* polymerizes an actin tail to propel it through the cytosol ^14, 15^, suggesting ER protrusions may form accidentally due to bacteria physically encountering the ER network. To investigate this hypothesis, we labeled A549-BiP-mNeonGreen-KDEL cells with the filamentous actin (F-actin) probe SIR-actin ^16^. We then quantified the percentage of ER-associated bacteria that had an actin tail. We found that most protrusive structures are associated with actin, whereas none of the vacuole-like ER structures had actin tails **(Fig. 1d, e)**. These results suggest that ER protrusive structures are likely a result of accidental collisions by actin-propelled bacteria.

We next investigated the vacuole-like ER structures, which appeared stably associated with bacteria **(Fig 1b)**. In uninfected cells, certain ER-mediated contact sites with other organelles last only several seconds ^13, 17, 18^, with a few lasting several minutes ^19, 20^. Using live-cell imaging, we quantified the duration of the vacuole-like ER interactions around bacteria over a 3 min period to determine if the ER forms stable interactions with *R. parkeri*. We observed that these structures were highly stable, with over 90% of them persisting for more than 2 min **(Fig 1f; Movie S1)**. Occasionally, we also observed these interactions spontaneously form within our 3 min time-lapse movies around a bacterium present in the cytoplasm (**Movie S2**). Our data show that, unlike *L. monocytogenes* or *S. flexneri*, a subpopulation of stationary *R. parkeri* forms a stable interaction with the ER.

### Immobile bacteria display increased interactions with the ER

The lack of actin tails associated with the vacuole-like ER structures prompted us to speculate that an *R. parkeri* mutant that cannot hijack the actin cytoskeleton would form more vacuole-like ER structures relative to WT *R. parkeri* (RpBFP). To test this, we used a mutant with a transposon insertion in *sca2*, a rickettsial gene that encodes a human formin mimic protein required for bacterial actin-based motility ^14, 15, 21^, and compared it to WT using live-cell imaging. Quantifying the rickettsia-ER interactions in aggregate across > 2000 bacteria per experiment showed that the percentage of *sca2*::Tn mutant bacteria interacting with the ER was nearly 25-fold higher than WT **(Fig. 2a)**. Additionally, the *sca2*::Tn mutant completely lacked any protrusive structures **(Fig. 2b)**, supporting our earlier conclusion that ER protrusions result from actin-propelled *R. parkeri*. Finally, these interactions were not observed with mutants of *L. monocytogenes* (Δ*actA*) and *S. flexneri* (Δ*icsA*) also impaired in actin-based motility ^22–25^ **(Fig. S1a, b)**. Together, these data show that a lack of actin-based motility increases the incidence of the unique interactions between rickettsia and the ER.

**Figure 2:**
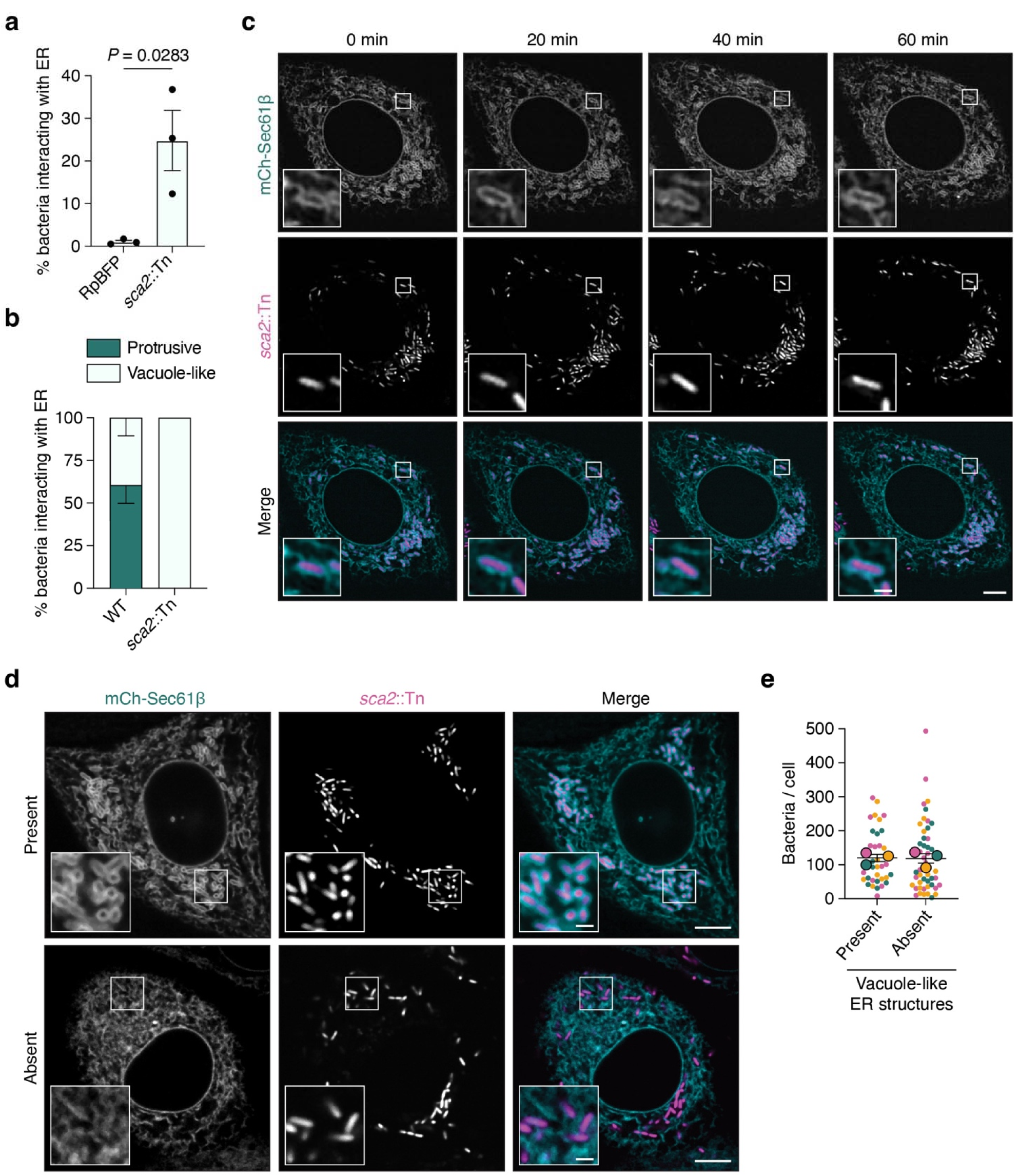
The *sca2*::Tn mutant displays increased interactions with the ER. **a**, Percentage of RpBFP or *sca2*::Tn mutant bacteria forming vacuole-like ER structures. RpBFP data are independent from that shown in Fig 1. Each dot represents the mean percentage of bacteria interacting with the ER for 3 independent experiments (> 4000 bacteria per independent experiment). Error bars represent SEM. Unpaired two-tailed *t*-test was performed. **b**, Relative frequency of RpBFP or *sca2*::Tn mutant bacteria forming protrusive or vacuole-like ER structures at 48 hpi. Error bars indicate SEM from 3 independent experiments (> 100 bacteria counted per independent experiment). **c**, Live-cell imaging snapshots at 48 hpi of *sca2*::Tn mutant bacteria (magenta) infecting infection A549-mCherry-Sec61β cells (cyan) captured at 5 min intervals for an hour. Scale bar, 5µm; inset 1µm. **d**, Live-cell imaging snapshots at 48 hpi of *sca2*::Tn mutant (magenta) infecting A549-mCherry-Sec61β cells (cyan). Images show cells with (present) or without (absent) vacuole-like ER structures. Scale bar, 5µm; inset scale bar, 1µm. **e**, Number of bacteria in cells that contain vacuole-like ER structures (present) or cells that lack them (absent) (images in d). The mean ± SEM of 3 independent experiments (outlined circles) are superimposed over the raw data from individual cells (unlined circles) (> 10 cells counted per independent experiment).

The interactions between the ER and *sca2*::Tn mutant bacteria were extremely stable, lasting more than 2 min **(Fig. S3)**. Because the *sca2*::Tn mutant interaction frequency was more abundant, we could track these interactions for longer and found that > 90% of the bacteria were stably associated with the ER for the entire hour of imaging **(Fig. 2c; Movie S3)**. Simultaneously, we observed that a subset of host cells displayed nearly uniform interactions between the *sca2::*Tn mutant and the ER, while other host cells were devoid of these interactions **(Fig. 2d)**. The bimodal nature of these interactions was not correlated with obvious differences in intracellular bacterial burdens, suggesting that these structures do not grossly impact bacterial replication or survival **(Fig. 2e)**. Thus, the *sca2*::Tn mutant strain emerged as a valuable model for further dissection of the rickettsia-ER interactions.

### *R. parkeri* is partially wrapped by the rough ER in multiple cell types

Stable interactions with the ER could occur by rickettsia gaining access to the ER lumen or from cytosolic bacteria being wrapped by the ER network. To differentiate between these possibilities, we used transmission electron microscopy (TEM) to resolve the rickettsia-ER interactions in A549 cells infected with the *sca2*::Tn mutant. We did not detect rickettsia inside the lumen of single-membrane structures. Instead, bacteria were partially wrapped by double membranes that contained ribosomes, indicating that the rough ER (rER) network wrapped around cytosolic bacteria **(Fig. 3a)**. We additionally observed similar structures in infected human microvascular endothelial cells (HMEC-1) **(Fig. 3b)**, a physiologically-relevant target of rickettsia species ^26–28^. Thus, the rickettsia-ER interaction is conserved in multiple cell types.

**Figure 3:**
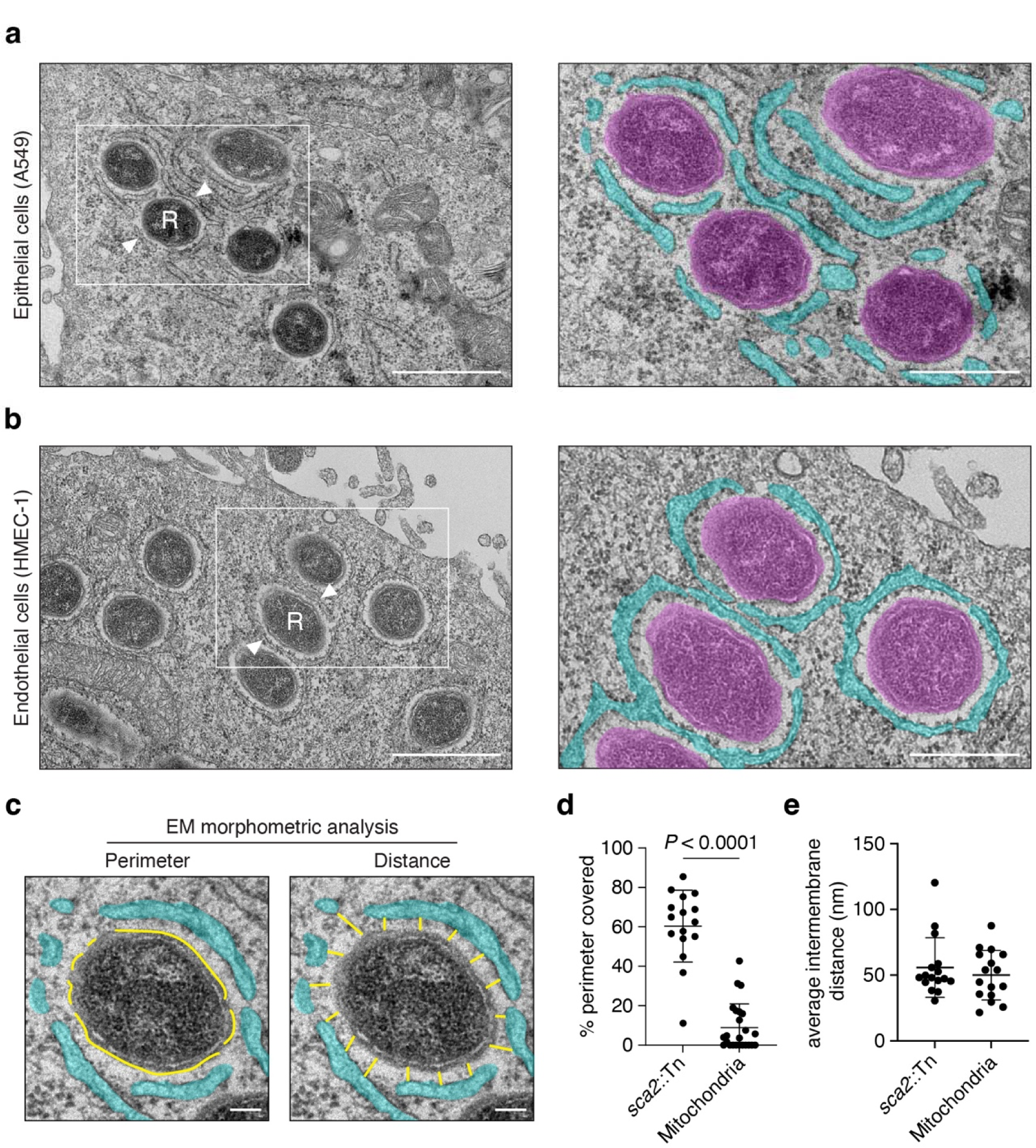
*R. parkeri* is partially wrapped by the rough ER in different cell types. **a** and **b**, TEM images at 30 hpi of A549 (a) and HMEC-1 (b) cells infected with *sca2*::Tn mutant bacteria. R, rickettsia; arrowhead, rER surrounding the bacterium. Insets to the right show bacteria in magenta and the ER in cyan. Scale bar, 1 µm; scale bar inset 500 nm. **c**, Examples of the 2D morphometric analysis used to quantify the average bacterial perimeter covered by the ER (left) or the average distance between ER and bacterial membranes (right) (from cells in a). ER, cyan; yellow lines represent traces measured. **d,** Percentage of the bacterial or mitochondrial perimeter covered by the ER (images in A). Each dot represents an individual bacterium or mitochondrion. (> 10 bacteria or mitochondria counted). Data shown are mean ± SD from an independent replicate. Unpaired two-tailed *t*-test was performed. **e,** Average distance between the rickettsial or mitochondrial membrane and the ER membrane. Each dot represents an individual bacterium or mitochondrion. (> 10 bacteria or mitochondria counted). Data shown are mean ± SD from an independent replicate. All data are a single representative replicate (n = 2).

### *R. parkeri* forms extensive membrane contact sites with the ER

Mitochondria-ER contact sites (i.e., MERCs) are 10–80 nm apart with variable percentages of the mitochondria perimeter wrapped by the ER (from 5 to close to 100%) ^7, 29–31^. Given the size similarities between mitochondria and rickettsia, coupled with the fact that both interact stably with the ER, **(Fig. 1f, 2d)** we hypothesized that rickettsia formed direct, interkingdom MCSs with the ER. Using 2D-morphometric analysis of our TEM data, we observed that approximately 58.7% of the bacterial perimeter was surrounded by the ER **(Fig. 3c, d)**. The extent of this interaction appeared to be specific because nearby mitochondria only showed interactions with the ER across approximately 8.4% of their perimeter **(Fig. 3c, d)**. We next measured the average distance between the ER membrane and the bacterial or mitochondrial surface along the perimeter of the contacts. We found that the ER had comparable average distances of approximately 55.7 nm and 50 nm to the rickettsial and mitochondrial surfaces, respectively **(Fig. 3c, e)**.

Another hallmark of an MCS is the presence of protein tethers that maintain the tight apposition of organelle membranes ^8, 32^. We predicted that similar tethers would bridge the ER and rickettsial membranes. To test this hypothesis, we used FIB-SEM to capture a greater extent of the rickettsial surface and attain a 3D view of the ER-rickettsia interactions. Using deep learning segmentation, we generated 3D models of the ER-rickettsia structures during infection (**Fig. 4a, Movie S4**). These 3D models showed that these ER-rickettsia associated structures are connected to the greater ER network **(Fig. 4a, Movie S5)** and confirmed our TEM data showing that > 50% of the rickettsial surface is covered by the ER **(Fig. 4b)**. In agreement with our hypothesis, we also observed regions of high-electron density that resembled tethering structures connecting the ER to the bacterial membrane **(Fig. 4c)**. These putative tethers exist throughout the ER-rickettsia interface and were present in all bacteria interacting with the ER. Collectively, these data indicate that *R. parkeri* forms MCSs with the ER, establishing the first example of a direct bacterium-ER contact site, or BERC.

**Figure 4:**
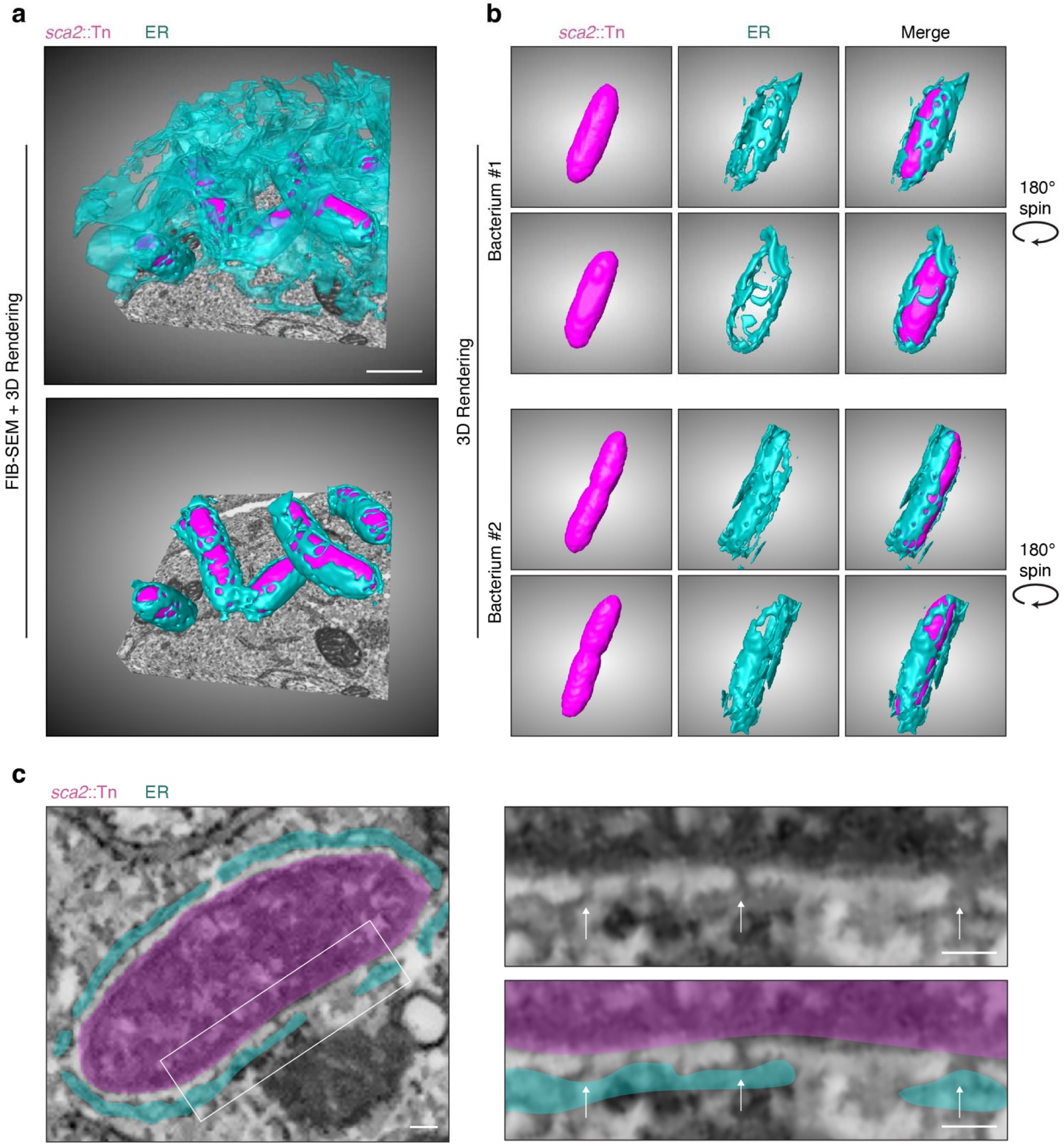
*R. parkeri* forms membrane contact sites with the ER. **a**, FIB-SEM image stacks and 3D rendering of ER-rickettsia structures in A549 cells infected with the *sca2*::Tn mutant (magenta) at 30 hpi. The top image shows a subvolume of the FIB-SEM image stack acquired with the entire ER network (cyan) and the *sca2*::Tn mutant displayed. The bottom image only shows 3D rendering of the ER membranes adjacent to the bacterial membranes. Scale bar, 1 µm. **b**, 3D rendering of individual bacteria surrounded by the ER from data shown in a. **c**, FIB-SEM slice of a *sca2*::Tn mutant bacterium (magenta) from a. Insets at right show electron dense regions (arrow) connecting the rickettsial (magenta) and ER (cyan) membranes. Scale bar, 100 nm.

### BERCs are reduced after VAPA and VAPB depletion

VAPs are ER-specific tethers that link the ER to different organelles ^33–38^. Indeed, loss of VAPA and VAPB expression decreases the frequency of ER-mediated MCSs ^34^. Consequently, we tested if VAPs also played a role in direct rickettsial-ER BERCs. We used RNA interference to deplete VAPA and VAPB in A549-mCh-Sec61β cells during infection with *R. parkeri* **(Fig. 5a)**. BERC frequency was then quantified using live-cell imaging after infection with the *sca2*::Tn mutant. Our results show that knockdown of *VAPA* and *VAPB* expression led to a partial but significant decrease in the frequency of host cells containing BERCs **(Fig. 5b)**, indicating that VAPs are important for BERC formation.

**Figure 5:**
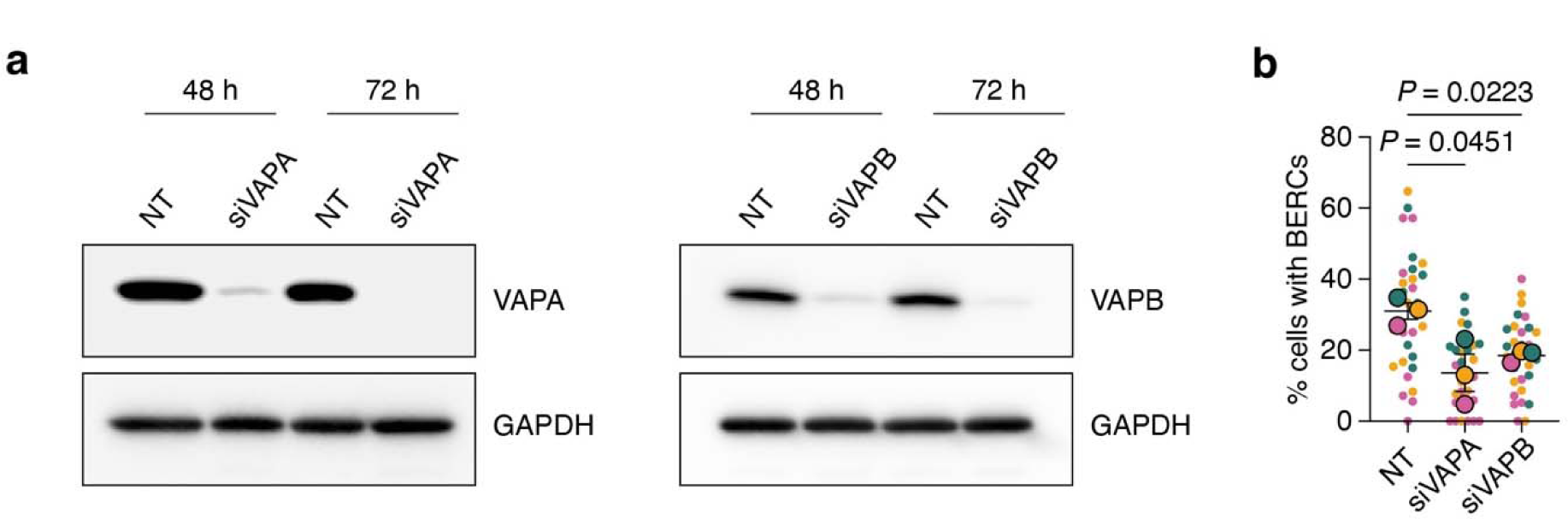
VAPA and VAPB depletion reduces the frequency of BERCs. **a**, Western blot analysis of A549-mCh-Sec61β cell lysates from b after indicated siRNA treatment at 48 h and 72 h post-transfection. GAPDH, loading control. **b**, Percentage of cells displaying BERCs at 24 hpi in A549-mCh-Sec61β cells infected with *sca2*::Tn mutant bacteria after the indicated siRNA treatment. The mean ± SEM of 3 independent experiments (outlined circles) are superimposed over the raw data representing the percent of cells with BERCs per FOV (unlined circles) (> 100 cells were quantified per condition per independent experiment). One-way ANOVA with Dunnett’s *post hoc* test was performed.

## DISCUSSION

The increased appreciation of interorganelle interactions in recent years has unveiled a variety of MCSs in eukaryotes ^39^. These closely apposed and tethered membranes facilitate numerous cellular processes, making them prime candidates for microbial control or mimicry ^9, 10^. Thus far, examples of pathogenic subversion of MCSs have been limited to interactions between eukaryotic membrane compartments ^4–6^. Indeed, the vacuolar bacterium *Chlamydia trachomatis* forms a contact site between its host-derived vacuole and the ER to maintain bacterial loads ^5, 6^ while the beta-herpesvirus human cytomegalovirus (HCMV) targets existing contact sites in the host to evade antiviral immunity ^4^. Here, we report the first example of a cytosolic bacterial pathogen forming a direct, interkingdom contact site with the ER during infection. We refer to these as bacterial-ER contact sites (BERCs) because the ER was directly tethered to the *R. parkeri* surface, resembling known mitochondrial-ER contact sites.

We observed an inverse relationship between actin-based motility and BERC formation, as the tail-less mutant *sca2*::Tn shows roughly 25-fold more interactions with the ER than WT *R. parkeri*. One of two scenarios could explain this observation: lacking actin-based motility increases the likelihood of a bacterium forming BERCs or decreases the chances of escaping BERCs once they are formed. Additionally, the bimodal phenotype of BERC formation observed with the *sca2*::Tn mutant indicates that motility is not the only layer of regulation. Understanding what role motility plays and what triggers BERC formation through host-or bacterial-specific mechanisms should elucidate if BERC formation is driven by a subset of bacteria or if all bacteria can form BERCs but do so at different phases of their life cycle.

The stability and average spacing of BERCs led us to speculate the existence of tether-stabilizing structures. Remarkably, we show that electron-dense regions of unknown identity connect the rickettsial and ER surfaces. These structures are similar to the molecular tethers at eukaryotic MCSs ^7, 40^ and our data suggest that the ER tethers VAPA and VAPB are involved. The incomplete loss of BERCs after VAPA or VAPB depletion could be due to trace amounts of protein left after siRNA treatment or could indicate the existence of other tethers linking *R. parkeri* to the ER network. At this time, we do not know what the VAPs are targeting because the *R. parkeri* surface lipidome and proteome is largely undefined. Further studies must be conducted to identify the bacterial factors that enable BERC formation. Once known, this discovery will facilitate functional dissection of the BERCs during infection.

It is unlikely that BERCs grossly impair bacterial survival through processes such as autophagy because BERCs were not correlated with lower bacterial burdens and the *R. parkeri* strains used in this study are resistant to autophagy ^41^. In addition to autophagy, interorganelle communication at MCSs regulates lipid metabolism, carbohydrate metabolism, apoptosis, inflammasome formation, and calcium signaling ^2, 3^. The obligate lifestyle of *R. parkeri* highlights that these pathogens are reliant on their host to survive and one or more of these MCS functions could be regulated by BERCs. For example, BERCs might promote metabolite exchange between the ER and the bacterium, and it will be important to investigate this possibility once our understanding of rickettsial metabolism improves. It is also possible that BERCs represent a unique adaptation to modulate host or bacterial function that remains to be identified.

Our discovery of a direct, interkingdom interaction between the bacterial pathogen *R. parkeri* and the ER challenges our traditional view of their cytosolic infectious life cycle. Indeed, this unique adaptation by *R. parkeri,* which was not found for other cytosolic pathogens we tested, exemplifies the exceptional strategies this pathogen has evolved and underscores the importance of better understanding its biology. It also remains to be seen if *R. parkeri* interacts with other organelle membranes, potentiating large-scale changes to the cellular landscape in favor of pathogen or host fitness. In the end, this work highlights an exciting new mode of host-pathogen interaction whose study may also deepen our understanding of MCS regulation and ER function.

## METHODS

### Cell lines

A549 (human lung epithelial), Vero (monkey kidney epithelial), and HEK293T (human embryonic kidney) cells were obtained from the University of California, Berkeley Cell Culture Facility (Berkeley, CA). HMEC-1 (human microvascular endothelial cell line-1) were a gift from Dr. Matthew Welch at the University of California (Berkeley, CA). A549 cells and HEK293T cells were maintained in Dulbecco’s modified Eagle’s medium (DMEM; Gibco, 11965118), supplemented with 10% fetal bovine serum (FBS). Vero cells were maintained in DMEM supplemented with 5% FBS. HMEC-1 cells were maintained in MCDB 131 liquid medium with no glutamine (Thermo Fisher, 10372019), supplemented with 10% FBS, 1 µg/mL hydrocortisone (Sigma-Aldrich, 50-23-7), 10 ng/mL epidermal growth factor (Corning, 354001), and 2 mM L-glutamine (Sigma-Aldrich, 56-85-9).

A549 cells expressing mCherry-Sec61β or the BiP signal sequence (ss) fused to mNeonGreen were generated using lentiviral transduction, as previously described ^42^. Briefly, the mCherry-Sec61β fusion was subcloned from mCh-Sec61β (Addgene, 49155) into the lentiviral plasmid FCW2IB-Lifeact-mWasabi ^42^ by replacing the LifeAct-mWasabi fusion, upstream of the IRES. For the BiP-mNeonGreen-KDEL fusion, the BiP signal sequence ^13^ and a KDEL sequence were fused the N- and C-termini of mNeonGreen, respectively. The BiP-mNeonGreen-KDEL fusion was then subcloned upstream of the IRES of FCW2IB-LifeAct-mWasabi by replacing the LifeAct-mWasabi fusion. Viral particles were packaged by transfecting HEK293T cells plated the day before at 2.5 × 10^5^ cells/mL (2 mL/well, 6-well plate) using calcium phosphate with 400 ng pMDL-RRE, 400 ng pCMV-VSVg, 400 ng RSV-Rev and 800 ng FCW2IB-mCherry-Sec61β (or FCW2IB-BiP-mNeonGreen-KDEL). Roughly 24 h after transfection, the media was replaced with 1.2 mL of fresh media to concentrate virus. 48 h after transfection, the concentrated supernatant was filtered through a 0.45 µm syringe filter and added to A549 cells at ∼ 2 × 10^5^ cells/mL. After 24 h of adding the filtered supernatant, cells were overlaid with fresh media containing 12 µg/mL of blasticidin (Sigma-Aldrich, 3513-03-9).

To achieve similar fluorescence levels, A549-mCherrySec61β cells were sorted based on mCherry fluorescence using the FACS Aria II Cell Sorter (BD Biosciences) at the Whitehead Institute for Biomedical Research Flow Cytometry Core Facility (Cambridge, MA). All sorted and parental cell lines were confirmed to be mycoplasma-negative in a MycoAlert PLUS assay (Lonza, LT07-710) performed by the Koch Institute High-Throughput Sciences Facility (Cambridge, MA).

### Bacteria plasmids and strains used in this study

RpBFP ^42, 43^, the *sca2*::Tn mutant ^15^ (a kind gift from Dr. Matthew Welch), and the *sca2*::Tn transformed with pRAM18dSGA-OmpApr-GFPuv (*sca2*::Tn pRAM GFP) ^42^ were generated as previously described. RpBFP refers to *R. parkeri* expressing two TagBFP cassettes cloned into pRAM18dRA ^42, 43^. All rickettsial strains were propagated at 33°C in Vero cells with DMEM containing 2% FBS, and isolated by mechanical disruption, as previously described ^42, 43^. Bacteria were aliquoted in brain heart infusion media (BHI; Fisher Scientific, DF0037-17-8) and stored at −80°C to decrease variability between replicates due to freeze-thaw cycles. Bacterial titers were determined as previously described via plaque assay ^42^.

The DasherGFP-expressing strain (LmDasher, strain PL1938 10403S:DasherGFP) was a kind gift from Dr. Erin Benanti (Aduro Biotech). To generate LmDasher, DasherGFP was codon-optimized for *L. monocytogenes* expression, cloned downstream of the *actA* promoter and integrated at the tRNA-Arg locus using pPL2. *L. monocytogenes* strains were grown in BHI. The GFP *L. monocytogenes* ActA deletion strain (Δ*actA*) was a gift from Dr. Michelle Reniere (University of Washington) and Dr. Dan Portnoy (UC Berkeley) and was generated by integrating pPL2-gfp ^44^ into the DP-L3078 (Δ*actA*) strain ^45^.

WT *S. flexneri* 2457T ^46^ and the isogenic IcsA mutant *icsA*::Ω (specR) ^47^ were gifts from Dr. Marcia Goldberg (Massachusetts General Hospital, Boston, MA). *S. flexneri* strains were grown on tryptic soy agar supplemented with 0.01% Congo Red (LabChem LC133507) and supplemented with 100 μg/uL spectinomycin (Sigma-Aldrich, 22189-32-8) or 100 μg/µL ampicilin (Sigma-Aldrich, 69-52-3) as needed. WT *S. flexneri* and 2457T *icsA*::Ω (specR) were transfected with the GFPmut3-expressing plasmid pFPV25.1 ^48, 49^ (Addgene, 20668) using electroporation, as previously described ^50^.

### Live microscopy of bacterial infections

All experiments were imaged with the Olympus IXplore Spin microscope system using an environmental chamber set to 5% CO_2_ and 37°C for *L. monocytogenes* and *S. flexneri* infections or 33°C for *R. parkeri* infections. Movies were acquired using an 100X UPlanSApo objective (1.35Lnumerical aperture) and Z Drift Compensation. Movies were deconvolved using Olympus CellSens software. All data quantification was performed in ImageJ, where bacteria were manually counted on the last frame of each movie.

To determine if *L. monocytogenes* forms interactions with the ER, 2 mL BHI cultures were inoculated with *L. monocytogenes* and grown for 16–18 h overnight at 30°C without shaking. A549-mCh-Sec61β cells at 2.5 cells/cm^2^ were grown in 20 mm Mattek dishes (MatTek Corporation, P35G-1.5-20-C). Bacteria from 0.5 mL of culture grown overnight were pelleted by centrifugation at 10,000L×Lg for 30 s at room temperature and resuspended in 500 µL PBS (Phosphate-buffered saline). Resuspended bacteria (LmDasher or Δ*actA*) were used to infect cells by adding suspension directly to cells at an MOI of 1–2. Infections were carried out at 37°C for at least 4 h before imaging. Before imaging, the dishes were washed two times and overlayed with FluoroBrite DMEM (Thermo Fisher, A1896702) containing 10% FBS. Movies were acquired at 20 s intervals for at least three minutes. Infections with both strains were imaged and > 30 cells representing > 1000 total bacteria were imaged per strain for each replicate (n = 3).

To determine if *S. flexneri* forms interactions with the ER, *S. flexneri* was grown overnight in tryptic soy broth (TSB) at 37°C with shaking. Overnight cultures were subcultured 1:200 and grown at 37°C with shaking for 2–3 h until cultures reached exponential phase (OD600 ≈ 0.4–0.6). Bacteria from 1 mL of culture were pelleted by centrifugation and resuspended in 250 μL PBS. Resuspended bacteria were used to infect A549-mCh-Sec61β cells at 0.6 × 10^5^ cells/cm^2^ in an 8-well chambered coverglass with a No. 1.5 borosilicate glass bottom (Thermo Scientific, 155360) at an MOI of 100. Infected cells were incubated for 30 minutes at 37°C and 5% CO_2_. Cells were washed twice with warm PBS and media was replaced with FluoroBrite DMEM containing 10% FBS and 25 µg/mL gentamicin. Cells were incubated an additional 1–1.5 hours until intracellular bacteria were clearly visible. Movie acquisition and processing was done as with *L. monocytogenes* infections.

To determine the frequency of ER-rickettsia interactions, A549-mCh-Sec61β cells on 8-well chambered coverglass (at 0.6 × 10^5^ cells/cm^2^) were infected with RpBFP (WT) or *sca2*::Tn (*sca2*::Tn pRAM-GFP) at an MOI of 0.5–2 for 28 and 48 h at 33°C (note, *R. parkeri* requires a longer incubation to accommodate its slower replication rate relative to *L. monocytogenes* and *S. flexneri*). Before imaging, each well was washed two times and overlayed with FluoroBrite DMEM containing 10% FBS. Movies were collected at 20 s intervals for at least three minutes. Approximately 10 fields of view were imaged for each replicate (n = 3) and > 2000 bacteria were quantified.

To visualize actin dynamics in the context of ER-rickettsia interactions, A549-BiP-mNeonGreen-KDEL cells on 8-well chambered coverglass (at 0.6 × 10^5^ cells/cm^2^) were infected at an MOI of 0.5–1 with RpBFP and incubated at 33°C for 48 h. Before imaging, each well was washed two times with FluoroBrite DMEM containing 10% FBS. Then, cells were overlaid with SIR-Actin (Cytoskeleton, CY-SC001) at a final concentration of 1 µM in FluoroBrite DMEM containing 10% FBS and incubated for 1 h. Imaging was done as described above for live cell imaging of rickettsial infections. Approximately 10 fields of view were imaged per strain for each replicate (n = 2) and > 100 bacteria interacting with the ER were quantified.

To determine the stability of ER interactions with the *sca2*::Tn mutant over a 1 h period, A549-mCh-Sec61β cells (2.5 × 10^5^ cells/cm^2^) in an 8-well chambered coverglass were infected by centrifuging cells at 200L×Lg for 5 min at room temperature with an MOI of 1.5–2. Infected cells were incubated at 33°C for 24 h, and imaging proceeded as above for rickettsial infections. Approximately 10 infected cells displaying ER interactions with the *sca2*::Tn mutant were imaged per replicate (n = 3). Movies were captured at 5 min intervals and ER interactions were tracked over the 1 h period. > 20 cells were quantified per replicate.

### Transmission Electron Microscopy

A549 and HMEC-1 cells seeded at approximately 3.0 × 10^5^ cells/cm^2^ or 1.5 × 10^5^ cells/cm^2^, respectively, in a 10cm^2^ dish were infected with the *sca2*::Tn mutant at an MOI of 0.5–1. Briefly, media was aspirated from cells, and cells were washed one time with PBS and overlaid with 1 mL of BHI covering the entire layer of cells. Then, the bacteria were added to cells and incubated via slow rocking at 37°C for 30 min. After, infected cells were overlaid with DMEM containing 10% FBS (A549) or MCDB 131 liquid medium without glutamine, supplemented as described above (HMEC-1) and incubated at 33°C for 30 h. Cells were then fixed using Karnovsky Fixative Reagent Grade (3% glutaraldehyde and 2% formaldehyde in 0.1 M phosphate buffer, pH 7.4) (Electron Microscopy Sciences, 15732-10) for 1 h at room temperature. After incubation, cells were scraped and centrifuged at 10,000L×Lg for 5 min at room temperature in 50 mL conicals. The pellet was transferred to a 1.5 mL Eppendorf, where it was centrifuged again at 10,000L×Lg for 2 min at room temperature and stored in fresh Karnovsky Fixative Reagent at 4°C until embedding.

Fixed cells were post fixed in 1% osmium tethaoxide in veronal-acetate buffer. The cells were stained en block overnight with 0.5% uranyl acetate in veronal-acetate buffer (pH 6.0), then dehydrated and embedded in Embed-812 resin. Sections were cut on a Leica Ultra Microtome with a diatome diamond knife at a thickness setting of 50 nm, stained with 2% uranyl acetate, and lead citrate. The sections were examined using a Hitachi 7800 TEM 80KV and photographed with an AMT CCD camera. Sample preparation after fixation and imaging was performed at the Harvard University Center for Nanoscale Systems (CNS) (Cambridge and Allston, MA). Two replicates were performed per cell line.

2D morphometric analysis to determine the percent bacterial or mitochondrial perimeter covered by the ER as well as the average distance between the ER and bacterial or mitochondrial membrane was performed in A549 cells by tracing the subcellular structures in Image J. To determine the average distance between the ER and bacterial or mitochondrial membrane, distances were quantified every 100 nm for ER membranes that were separated ≤ 130 nm from the surface of either bacteria or mitochondria. To determine the percent perimeter covered, the bacterial and mitochondrial surface that was ≤ 130 nm from an ER membrane was traced. Approximately > 10 bacteria and > 10 mitochondria were quantified per replicate (n = 2).

### Focus Ion Beam Electron Microscopy (FIB-SEM)

To locate an infected cell for 3D FIB-SEM imaging, a thin plastic section was lifted off the block face using a microtrome knife, transferred onto a TEM grid and scanned in a TEM for the presence of bacteria (Leica Ultramicrotome, Hitachi 7800 TEM). To reduce the search area and facilitate determining the orientation of the thin section, the block had been trimmed prior to a trapezoidal cross section with an approximate edge length of 500 µm. Once a cell with the desired ultrastructure was identified, a series of images were recorded at cascading magnifications keeping the region of interest in the image center. The magnification was lowered until a block edge or corner was visible in the image. This approach made it possible to start the search in the SEM on an easily recognizable region of the block and use characteristic features at varying length scales in the corresponding back-scatter image to trace a path back to the actual region of interest.

FIB-SEM data were acquired on a Zeiss Crossbeam 550 using the Atlas software package. The cell was prepared for automatic serial sectioning by first covering its visible region with a 1-µm thick Pt film via ion-beam assisted deposition. A series of line markings carved into the film with the ion beam act as fiducial markers for drift correction and the measurement of the progression of the ion beam during the run. The grooves are subsequently covered with a 1-µm thick carbon film to reduce curtaining artifacts, generate contrast, and ultimately preserve the markings throughout the run. After clearing material in front of the region of interest to a visual depth of 20 µm, data were recorded using the energy-selective back-scatter detector with the electron beam operating at 1.5 kV and 2nA. Cross section images with a 5 nm pixel size and a dwell time of 10 µs were recorded in 10 nm slice intervals. Milling was performed with the ion beam set to 30 kV and a current of 700 pA.

Images were aligned via cross-correlation and denoised with the non-local means filter using an in-house Matlab-based software package. Due to the relatively long dwell time, images contained small distortions along the slow scan direction. A stripe-based cross correlation approach was designed to carefully measure displacements within stripes of consecutive images which allowed to construct a displacement field for each image that is subsequently used to correct for locally varying drift and to ensure exact alignment throughout the volume of the dataset. Analysis scripts are available upon request through the corresponding author. The final data volume of 15 × 15 × 15 µm^3^ ultimately contained 17 bacteria enveloped by the cell ER. Deep learning data segmentation was performed using the Object Research Systems (ORS) Dragonfly software to extract the organelles of interest. A U-Net model with five depth levels, a starting number of 64 convolutional kernels and a patch size of 64 pixels was successfully trained using 3 manually segmented image slices. The model was then applied to the rest of the FIB-SEM image stack. 3D models were generated using the segmentation data and the 3D visualizer in ORS Dragonfly.

### VAPA and VAPB depletion

To determine the frequency of cells containing BERCs after VAPA or VAPB depletion, A549-mCh-Sec61β at approximately 0.6 × 10^5^ cells/cm^2^ in an 8-well chambered coverglass were transfected with 10 nM of pooled siRNA against *VAPA* (Horizon Discovery, L-021382-00-0005), *VAPB* (Horizon Discovery, L-017795-00-0005) or a non-target control (Horizon Discovery, D-001810-10-05) using Lipofectamine™ RNAiMAX Transfection Reagent (ThermoFisher, 13778030). Cells were grown for 48 h at 37°C. At 48 h post-transfection, cells were infected at an MOI of 1.5–2 with the *sca2*::Tn mutant by centrifuging cells at 200L×Lg for 5 min at room temperature. The infected cells were then incubated at 33°C for 24 h. Imaging was performed as described above, with approximately 10 fields of view imaged per condition for each replicate (n = 3) and > 300 cells quantified. Data quantification was manually performed on ImageJ where cells containing BERCs were quantified per FOV.

### Western Blot

To confirm siRNA-mediated silencing of *VAPA* and *VAPB* expression, transfections were completed as above except cells were plated in 24-well dishes. Cells were collected at 48 and 72 h and lysed on ice for 10 min in immunoprecipitation lysis buffer (50 mM HEPES, 150 mM NaCl, 1 mM EDTA, 10% glycerol, 1% IGEPAL). The cell debris was then cleared via centrifugation at 16,100 × g 4°C for 10 min. Lysates were analyzed by Western blotting using rabbit anti-VAPA (Proteintech, 15275-1-AP) and mouse anti-VAPB (Proteintech, 66191-1-Ig) and rabbit anti-GAPDH (Cell Signaling, 2118S).

### Statistics

The statistical analysis and significance are noted in the figure legends for each respective graph. Data are determined to be statistically significant when p < 0.05 by unpaired t-test (two-tailed) and one-way ANOVA with post hoc Dunnett’s test. Statistical analyses were performed using GraphPad PRISM 9.

## ACKNOWLEDGEMENTS

We thank Matthew Welch, Erin Benanti, Michelle Reniere, Dan Portnoy, and Marcia Goldberg for sharing strains and reagents. We thank Jon McGinn, Iain Cheeseman, Adam Martin, and Allison Scott for providing critical feedback when reading our manuscript. We thank Nicki Watson for help preparing and imaging the TEM samples, as well as for providing training in imaging on the TEM. This work was performed in part at the Harvard University Center for Nanoscale Systems (CNS); a member of the National Nanotechnology Coordinated Infrastructure Network (NNCI), which is supported by the National Science Foundation under NSF award no. ECCS-2025158. This work was also performed in part at the Flow Cytometry Core Facility at the Whitehead Institute for Biomedical Research. This work was supported in part by the National Institutes of Health (NIH) Grants no. T32GM-007287 (YAS) and R01GM-141025 (to RLL).

## AUTHOR CONTRIBUTIONS

YAS and RLL conceived and designed the study. YAS performed the experiments unless otherwise stated. PJW performed the live microscopy *S. flexneri* experiments. SK processed the FIB-SEM samples and completed the deep learning segmentation. YAS and RLL wrote the manuscript with input from all the authors.

## MATERIALS & CORRESPONDENCE

Further information and requests for reagents may be directed to and will be fulfilled by Rebecca Lamason, (RLL).

## DATA AVAILABILITY

The data generated and analyzed in this study are available from the corresponding author upon request.

---

**Supplemental Figure 1:**
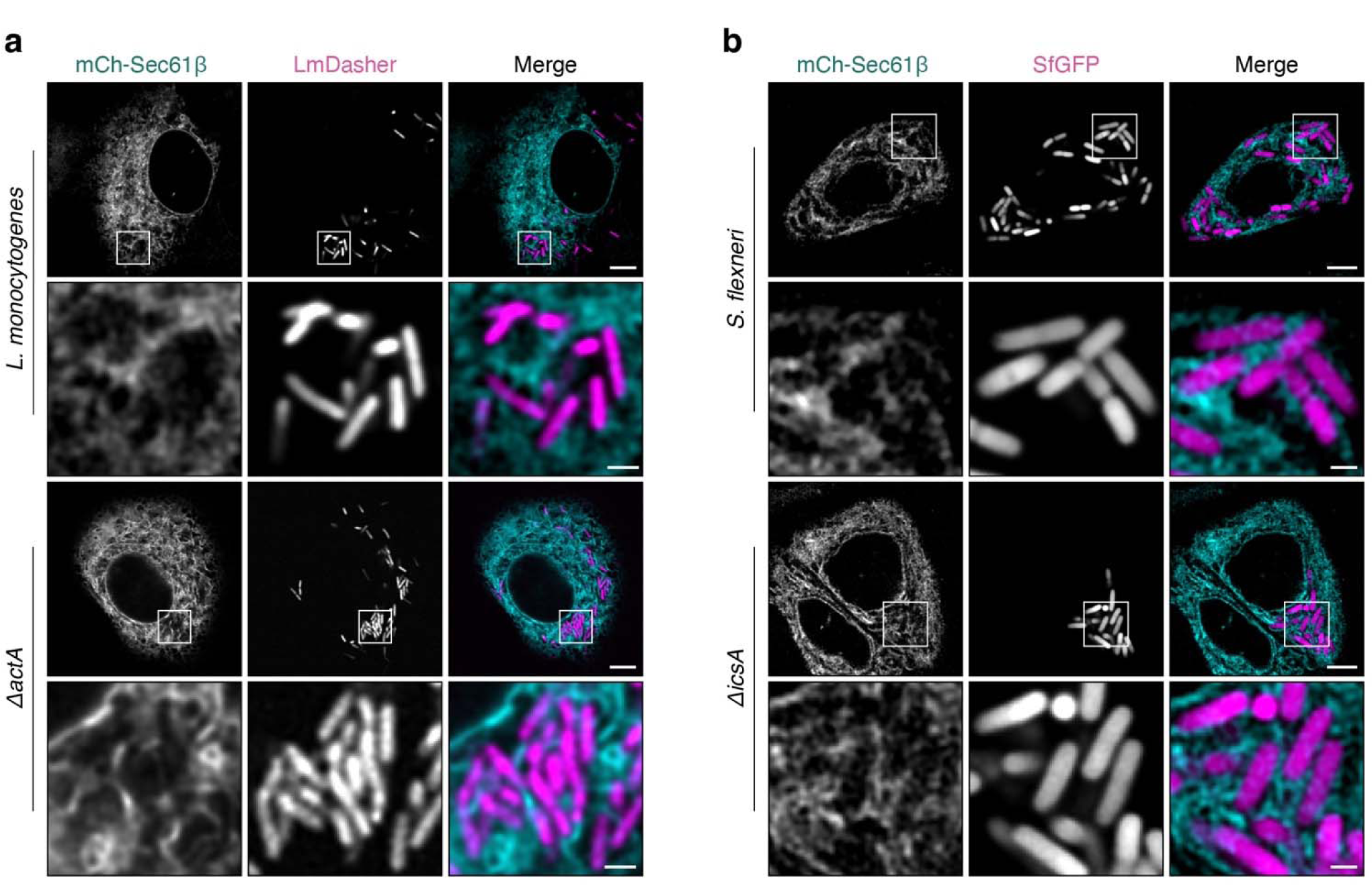
Stable ER interactions are not formed by *L. monocytogenes* or *S. flexneri*. **a**, Live-cell imaging snapshots at 4 hpi of A549-mCh-Sec61β cells (cyan) infected with DasherGFP-expressing WT or GFP-expressing Δ*actA* mutant strains of *L. monocytogenes* (magenta). Scale bar, 5µm; inset scale bar, 1µm. **b**, Live-cell imaging snapshots at 2 hpi of A549-mCh-Sec61β cells (cyan) infected with GFP-expressing WT or Δ*icsA* mutant strains of *S. flexneri* (magenta). Scale bar, 5µm; inset scale bar, 1µm. Data are representative of 3 independent experiments.

**Supplemental Figure 2:**
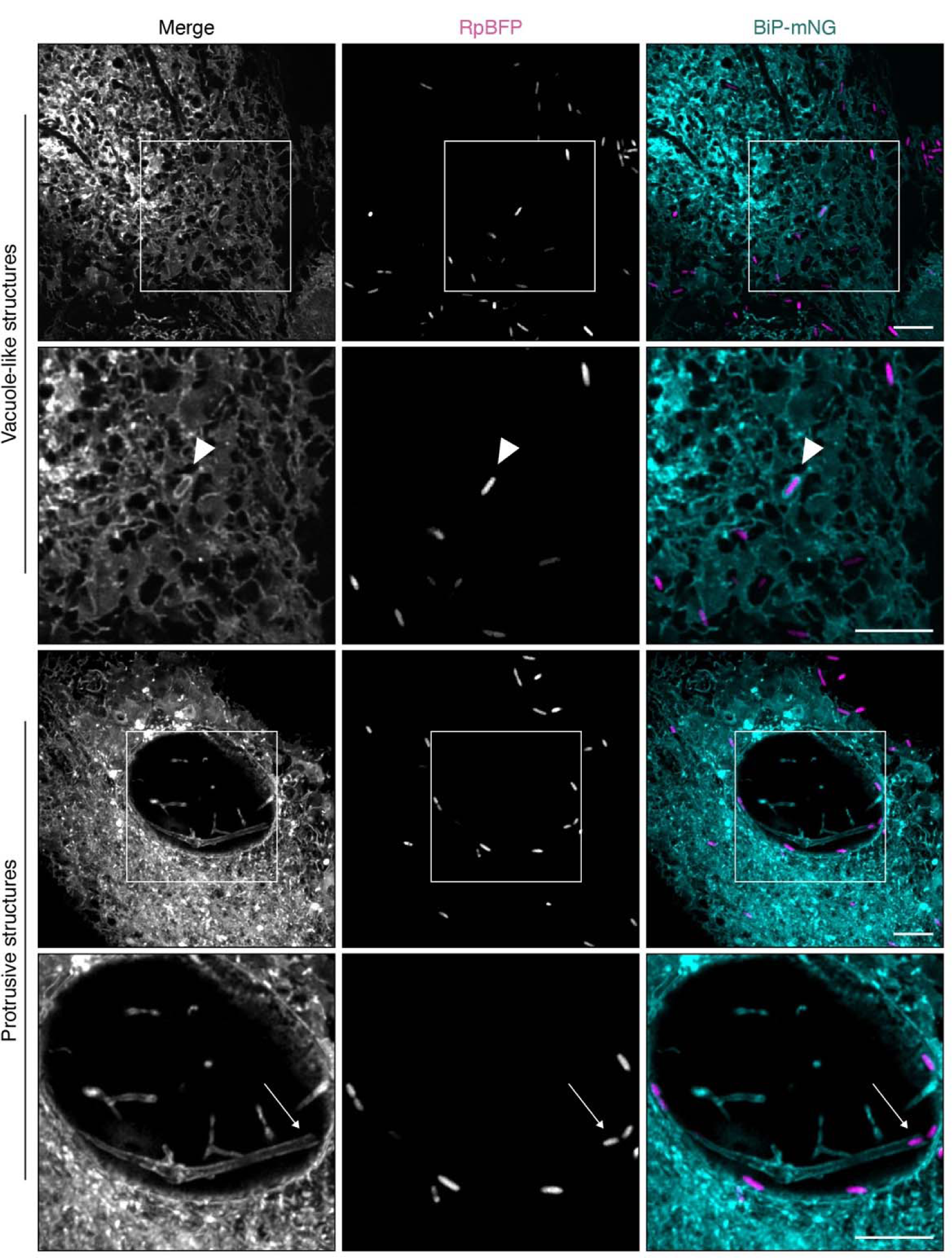
*R. parkeri* interacts with the ER. Live-cell imaging snapshots of A549 cells expressing an ER marker (BiP-mNeonGreen-KDEL, cyan) infected with *R. parkeri* expressing BFP (magenta) at 28 hpi. Arrowhead in inset points to *R. parkeri* in vacuole-like ER structure. Arrows in the inset point to *R. parkeri* in protrusive structures. Scale bars, 5µm.

**Supplemental Figure 3:**
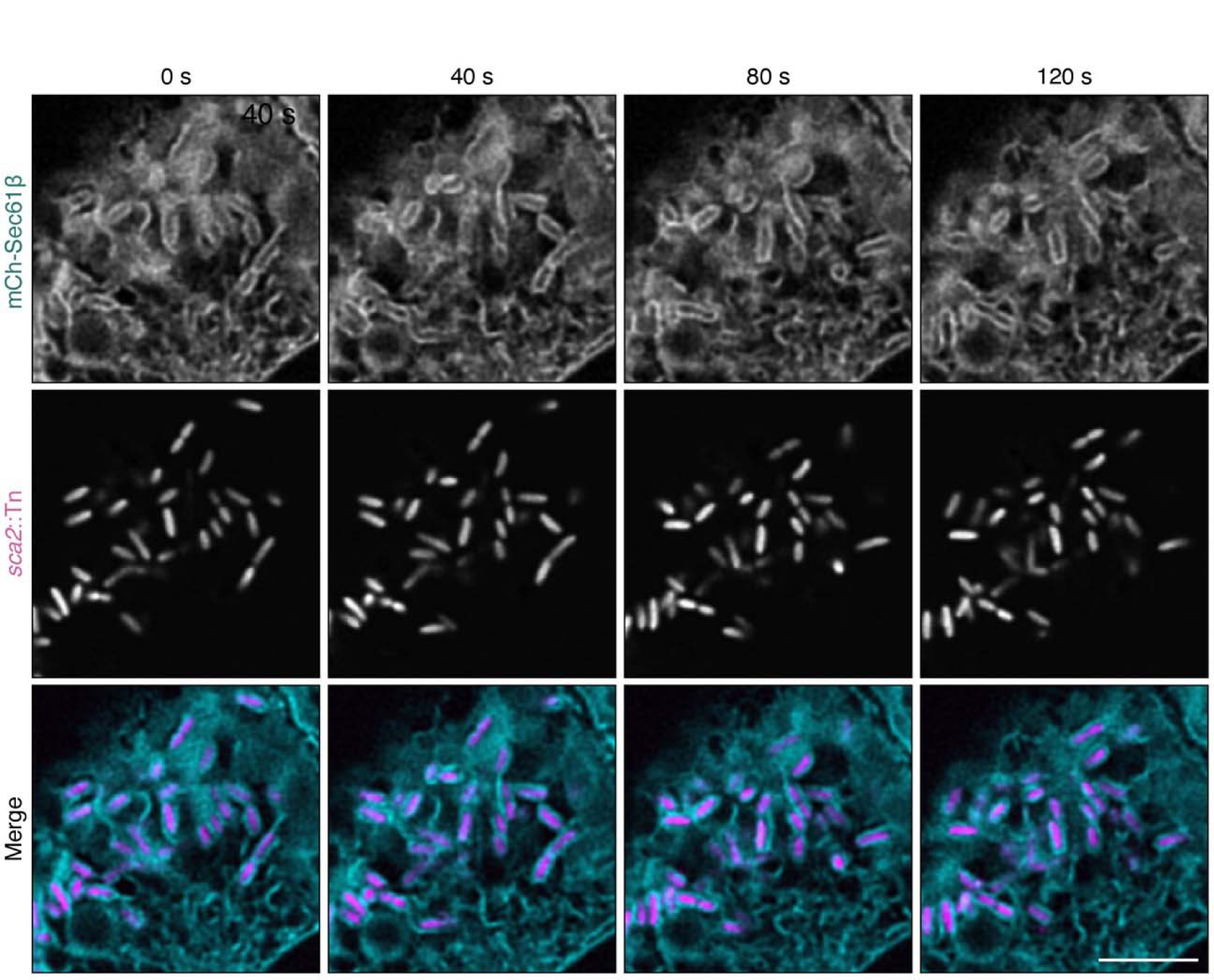
The *sca2*::Tn mutant forms stable interactions with the ER. Live-cell imaging snapshots at 48 hpi of A549-mCh-Sec61β cells (cyan) infected with the *sca2*::Tn mutant (magenta) forming ER vacuole-like structures. Images show the same bacterium during a 3-min time-lapse movie with frames captured every 20 s, similar to data acquired for WT in Fig. 1f.

**Supplemental Movie 1: *R. parkeri* forms stable interactions with the ER**

A549-mCh-Sec61β cells (cyan) infected with RpBFP (magenta) for 28 h show stable vacuole-like ER structures around the bacterium (arrow) (see also Fig. 1f). A single z slice is shown and frames were captured every 20 s. Timestamp shows seconds (s). Scale bar, 2.5 µm.

**Supplemental Movie 2: Formation of a vacuole-like ER interaction around *R. parkeri***

A549-BiP-mNeonGreen-KDEL cells (cyan) infected with RpBFP (magenta) for 28 h showing the formation of a vacuole-like ER structure around a bacterium (arrow). A single z slice is shown and frames were captured every 20 s. Timestamp shows seconds (s). Scale bar, 2.5 µm.

**Supplemental Movie 3: The *sca2*::Tn mutant forms long-lived interactions with the ER**

A549-mCh-Sec61β cells (cyan) infected with the *sca2*::Tn mutant (magenta) for 24 h show very stable vacuole-like ER structures around the bacterium. A single z slice is shown and frames were captured every 5 min. Timestamp shows minutes (min). Scale bar, 2.5 µm.

